# Analysis of *Escherichia coli* Flagellin Gene Expression in Simulated Spaceflight Growth Conditions

**DOI:** 10.1101/274621

**Authors:** Kyle M. Hetrick, Elizabeth A. Juergensmeyer, Jeffrey O. Henderson

## Abstract

Population growth curves of *Escherichia coli* (*E. coli*) cultures propagated in spaceflight environments demonstrate an extended log phase duration and an increased final cell concentration compared to *E. coli* cultures grown on Earth. We suggest that during spaceflight, a lack of convective mixing in the growth medium alters a cell’s microenvironment such that the cell at first devotes its resources primarily to motility and later shifts most of its resources to reproduction. As flagellin is the major protein component of flagella, the motility structure of *E. coli*, we propose that the expression of the flagellin gene (*fliC*) can be used as a diagnostic marker for determining if the bacterium is in fact altering its allocation of cellular resources to motility during specific growth phases. We have used semiquantitative RT-PCR to analyze the expression of *fliC* in cultures grown in laboratory apparatus that simulate various aspects of the spaceflight environment. The data from this pilot study indicate that *fliC* expression increases initially under stationary, slow clinorotation, and fast clinorotation growth conditions, peaking at mid-exponential phase and decreasing dramatically in stationary phase. These data support the hypothesis that metabolic resources are being directed towards forming the flagellum, the bacteria motile structure, before shifting resources to growth, reproduction, and maintenance.

## Introduction

In the space flight (microgravity) environment, bacteria show altered growth patterns compared to controls grown simultaneously on the ground. In minimal media, the bacterium *Escherichia coli* (*E. coli*) strain K12, substrain MG1667, has been shown to grow more rapidly, and ultimately achieve a higher final cell density (Reviewed in Juergensmeyer *et al.*, 2007; Klaus *et al*., 1997, 2004) in microgravity compared to Earth (standard gravity; 1 *g*). One hypothesis for rapid growth in microgravity is lack of buoyancy-driven convection currents, leading to minimal flow within a column, resulting in a microenvironment around each cell in which nutrients are not carried to the cell, nor are waste products carried away. After a period of cell growth, the environment within the culture column contains pockets of nutrient-rich and nutrient-poor areas (Klaus *et al*., 1997, 2004).

Motility in bacteria is not similar to motility in metazoans, in that the effort put forth by bacterial cells to be motile depends heavily on the amount of nutrients present. We propose the “pocket hypothesis” as a model for bacterial motility in the spaceflight (microgravity) environment. In a low-nutrient environment, a bacterial cell is almost constantly motile, with occasional pauses to evaluate the environment. The cell is constantly producing flagellin, the protein used to make the flagellar filament, the bacterial structure for motility (Zhou *et al*., 2015). Therefore, an analysis of flagellin gene (*fliC*) expression may indicate changes in the amount of resources that cultures of motile *E. coli* are devoting to motility as the population expands and finally plateaus. However, in a nutrient-rich environment, the cell can minimize motility, with the dual advantages of conserving the energy required directly for motility, and also the energy required to produce flagellin. Thus, a cell in a nutrient-rich pocket would not only have more catabolites available, it would also be able to decrease metabolic expenditure on motility and use that metabolic energy for growth and reproduction. A cell in a nutrient-poor environment would conversely spend more metabolic energy on motility and less on growth and reproduction (Wadhams and Armitage, 2004). A cell in a well-mixed environment would attain a median value of metabolic expenditure, with some portion of its metabolic process spent on motility, and the remainder on growth and reproduction. Therefore, the “pocket” environment hypothesized to exist during spaceflight (Klaus *et al.*, 1997, 2004) may allow increased growth efforts by inducing bacterial cells to first produce flagella, become motile, and then, when in a nutrient-rich area, allow the cells to store large amounts of metabolites and reproduce at an increased rate, without expending metabolites on a motility apparatus that already exists. When the nutrient “pocket” is depleted, the cells are not required to produce additional flagellin, but may move utilizing stored energy and previously synthesized flagella.

Due to the rarity and expense of spaceflight experiments, very few have been done. Neither the proximal nor the ultimate causes of the growth curve changes in microgravity are known (Reviewed in Klaus *et al*. 1997). However, there are multiple ground devices that are capable of mimicking one or more aspects of space flight. Clinorotation (Fig. 1) rotates a sample vertically around a central point, resulting in a randomized gravity vector. Gravity is at all times pulling on the sample, but because of the rotation, the organism perceives the gravity as coming from a constantly changing source. When a column of liquid is clinorotated, convection is reduced due to the constant change in direction of buoyant flow (Kacena *et al.*, 1999a,b).

**Figure 1.**
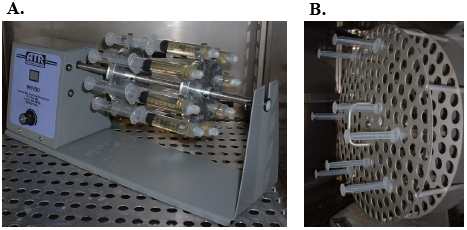
Simulation of randomized gravity vectors. (A) Fast clinostat holding hard-sided growth chambers. (B) Slow clinostat holding hard-sided growth chambers. Clinostats simulate the randomized gravity vectors that bacteria experience in a spacecraft undergoing a continuous state of free-fall around the Earth (although in an orbiting spacecraft, the magnitude of the gravity vectors would be much less). Convection is greatly reduced in clinorotated vessels.

Another, less well known, aspect of the spaceflight environment is vibrational acceleration. When gravitational forces are removed or reduced, the dominant accelerative force on an organism is vibration. On Earth, vibration is considered negligible, and often ignored, but when gravity is reduced, the vibration perceived from a motor, compressor, shaker, or other device is proportionally much larger (Nelson and Jules, 2004). Vibration of *E. coli* at 60 rpm (1 Hz; Fig. 2), on Earth, results in the same changes in the growth curve seen in space-grown bacteria (Juergensmeyer *et al*., 2007).

**Figure 2.**
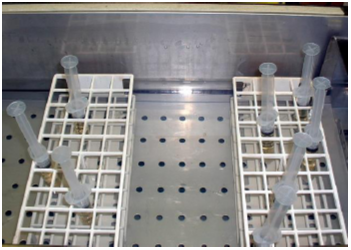
Simulation of vibrational forces encountered in spaceflight. A shaking water bath simulates the vibrational accelerations to which the bacteria would be exposed in the spaceflight environment. In microgravity, vibration may be the major accelerational force acting upon the organisms. The 10-mL syringes approximate the Fluids Processing Apparatus (BioServe Space Technologies, Boulder, CO), the “test tubes” of biological experiments performed in space.

Analyzing the growth of bacteria, especially pathogenic strains, is important and necessary for providing effective treatment of astronauts in the spaceflight environment. Using the non-pathogenic strain, *E. coli*, will present insights into possible mechanisms of increased bacterial growth in the microgravity environment associated with spaceflight. Analyzing *fliC* expression in *E. coli* grown under simulated spaceflight conditions may generate insight as to how motile bacteria grown in a spaceflight environment alter their distribution of metabolic resources over time.

## Experimental Procedures

### Cell Culture

A turbid overnight culture of *E. coli* strain K-12, substrain MG1667 (ATCC 10798), a motile strain, was used to inoculate fresh Luria-Bertani broth (LB; Becton, Dickinson and Company; Franklin Lakes, NJ, USA) at a ratio of 1 mL to 500 mL LB. The freshly inoculated broth was used to fill Luer-Lok 10-mL syringes (hard-sided tube; Klaus *et al*., 1997) and subjected to linear vibration (60 rpm = 1 Hz; see Juergensmeyer *et al*., 2007) in a shaker water bath (Precision Scientific; Winchester, PA or VWR; Scientific Products, West Chester, PA, USA), slow clinorotation (0.25 rpm), fast clinorotation (80 rpm), or stationary growth conditions at 37°C in an incubator. Optical density at 600 nm was monitored throughout culture growth. Bacteria were collected at lag, lag/exponential, mid-exponential, and stationary phases into RNA*later* (Ambion; Austin, TX, USA) and stored at –80°C.

### Semiquantitative Reverse Transcription-Polymerase Chain Reaction Amplification Analysis of fliC

Total RNA from bacterial pellets was purified using TRIZOL as described by the manufacturer (Invitrogen; Carlsbad, CA, USA). A semi-quantitative RT-PCR methodology was used as previously validated (Giannoni *et al*., 1994, 1995; Henderson *et al*., 2001). In brief, 20 μg of RNA was treated with 2 units RQ1 DNase at 37°C for 1 h in 100 μL 40 mM Tris-HCl pH 7.6, 6 mM MgCl2, 10 mM NaCl, 40 U RNasin (Promega; Madison, WI, USA). The RNA was sequentially extracted with phenol:chloroform and chloroform, precipitated with ethanol at –80°C, washed once with 75% ethanol and resuspended in RNase-, DNase-, protease-free water (Gibco BRL). RNA concentration was confirmed by measuring the absorbance at 260 nm. Five nanograms of RNA were reverse-transcribed using the SuperScript III First-Strand Synthesis System in the presence of 200 ng random hexamer (dN6) as described by the manufacturer (Invitrogen). The *fliC* cDNA was amplified using the primer pair *fliC-*F (sense; 5’-acagcctctcgctgatcact-3’) and fliC-R (antisense; 5’-gttatcatttgcgccaacct-3’). The housekeeping gene *rrlH*, which encodes 23S rRNA (Bulyk *et al.*, 2004), was used as the internal standard, and its cDNA was amplified using the primer pair rrlH-F (sense; 5’-cttaggcgtgtgactgcgta-3’) and *rrlH*-R (antisense; 5’-tgggttgtttccctcttcac-3’). The *fliC* and *rrlH* cDNA were amplified to the midpoint of linearity of the exponential phase using Platinum *Taq* PCR Mix (Invitrogen). PCR was performed for the established number of cycles in a Primus 96Plus thermal cycler (MWG Biotech AG; Ebersberg, Germany) equipped with a heated lid as follows: Initial heat denaturation at 94°C for 2 min, then 94°C for 30 sec, annealing at 55°C for 30 sec, and extension at 72°C for 1 min. A final 10 min extension was added after the last cycle. Amplicons were purified using the QIAquick PCR Purification Kit (Qiagen; Valencia, CA, USA) from agarose gels, and DNA concentrations determined at 260 nm in a BioPhotometer (Eppendorf AG; Hamburg, Germany) using ultraviolet light transparent UVett cuvettes (Eppendorf AG). Under the conditions detailed above, the midpoint of linearity of the exponential phase of *fliC* and *rrlH* was determined to be 24 and 16, respectively (Fig. 3). The ratio of [DNA]*flg* to [DNA]*rrlH* was calculated for all samples.

**Figure 3.**
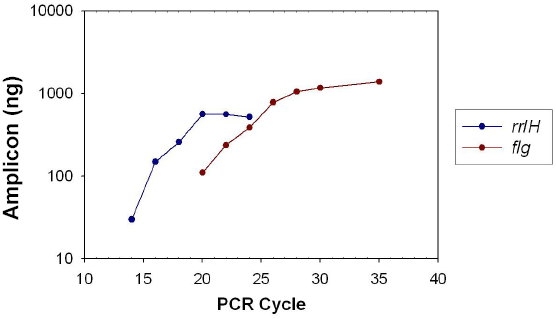
RT-PCR of *fliC* and *rrlH* to determine cycle number at mid-point of linearity of amplification. The graph shows the mean values of amplicon quantity obtained from three RT-PCR trials using RNA from *E.* coli grown under stationary conditions and collected at mid-log phase. Error bars are contained within each datapoint.

### Results and Discussion

We first compared the population growth dynamics of cultures grown under four different conditions: linear vibration, slow clionorotation, fast clinorotation, mimicking various aspects of growth in spaceflight, and static. Growth curves were generated (Fig. 4) and used to establish times corresponding to mid-lag, lag/exponential, mid-exponential, and stationary phases. Cultures grown under linear vibration, slow clinorotation, and fast clinorotation entered exponential phase before the control cultures (grown with no motion, static/stationary). However, only cultures grown under linear vibration reached a higher cell concentration by the end of the experiment; cultures grown under fast or slow clinorotation reached cell concentrations comparable to those of the control cultures, in agreement with previous studies (reviewed in Juergensmeyer *et al*., 2007).

**Figure 4.**
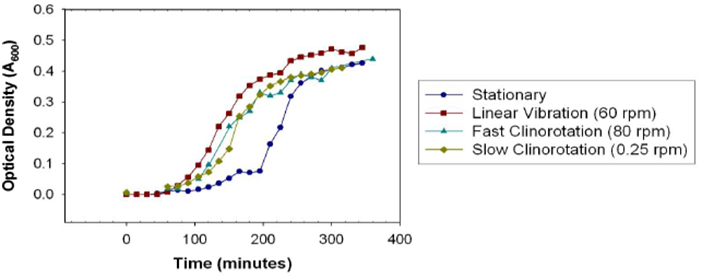
Culture growth during simulated spaceflight (microgravity) growth conditions. The data points show the mean OD600 of five independently grown cultures at each time point. Error bars are smaller than the data points.

We analyzed the *fliC* expression levels over time of cultures grown in simulated spaceflight growth conditions by isolating total RNA from independently grown cultures, synthesizing cDNA from the RNA, amplyfing the *fliC* cDNA, and normalizing the concentration of the *fliC* amplicon to that of the 23S rRNA housekeeping gene, *rrlH* (Fig. 5). These data indicate that cultures grown under slow or fast clinorotation behaved similarly to stationary cultures, with *fliC* expression greatest in mid-exponential phase. This was not the case for the cultures grown under linear vibration, where the data indicate that *fliC* expression were lowest in mid-exponential phase.

**Figure 5.**
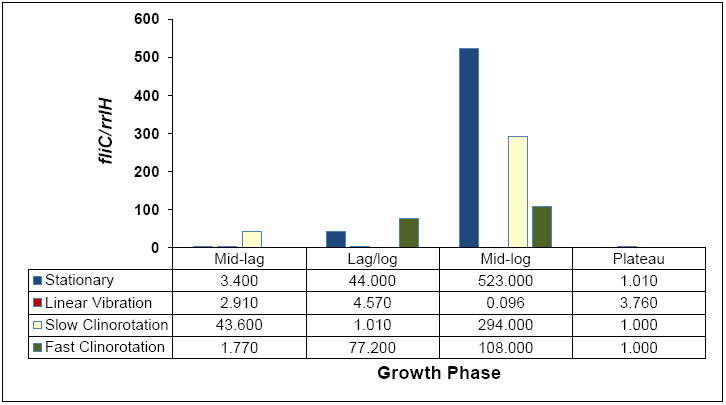
Semiquantitative RT-PCR of *fliC* mRNA. The bars show the mean ratio of *fliC* mRNA to the internal control, *rrlH* (*fliC/rrlH*), over two trials. Bacteria were harvested at the following growth points: middle of lag phase (Mid-lag), transition from lag phase to exponential phase (Lag/log), middle of exponential phase (Mid-log), and stationary phase (Plateau). Total RNA was isolated, converted to cDNA, amplified, and quantified as described in Experimental Procedures.

These data indicate that *fliC* expression increases initially (at the mid-lag and lag/exponential phases) under static, slow and fast clinorotation growth conditions, peaking at mid-exponential phase (mid-log) and decreasing dramatically in stationary phase (plateau). This trend in *fliC* expression is consistent with the prediction that *E. coli* in stationary and spaceflight environments would initially produce large amounts of flagellin to increase motility with a subsequent decrease in flagellin production over time as the flagella previously synthesized became sufficient for motility. This trend does not appear to hold for the linear vibration condition, suggesting that under mock spaceflight conditions, direction of motion makes a larger contribution to changes in *fliC* expression than speed of the motion.

To confirm these trends, the *fliC*/*rrlH* ratios will need to be determined for additional time points in triplicate. Future work would entail determining whether flagellin gene expression at the translational level correlates with mRNA levels with subsequent ascertainment of the molecular mechanisms involved in *fliC* regulation for *E. coli* grown under simulated spaceflight conditions.

## Acknowledgments

We thank Julie K. Henderson for editorial assistance. This work was carried out from 2004 to 2006 and supported by NASA Grant NAG2-1512 (EAJ, KMH), a Surbeck Summer Scholarship Grant from Judson University (2004; EAJ), a Summer Research Grant from Trinity International University (2004; JOH), and a one-semester sabbatical leave from Judson University (2017; JOH).

